# The low-density lipoprotein receptor-related protein-1 is essential for Dengue virus infection

**DOI:** 10.1101/2020.06.10.144089

**Authors:** Vivian Huerta, Alejandro M. Martin, Mónica Sarría, Osmany Guirola, Alexis Yero, Yassel Ramos, Dianne Pupo, Dayron Martin, Alessandro Marcello, Glay Chinea

## Abstract

Dengue virus (DENV) causes the most prevalent and rapidly spreading arboviral disease of humans. It enters human cells by receptor-mediated endocytosis. Numerous cell surface proteins have been proposed as DENV entry factors. Among these, the phosphatidylserine receptor TIM-1 is the only one known to mediate virus internalization. However, several cellular models lacking TIM-1 are permissive to DENV infection, suggesting that other receptors exist. Here we show that the Low-density lipoprotein receptor-related protein-1 (LRP1) binds DENV virions by interacting with the DIII of the viral envelope glycoprotein. DENV infection is effectively inhibited by the purified receptor at 5×10^−8^ mol/L and the interaction of the envelope protein with LRP1 is also blocked by a natural ligand of LRP1. Depletion of LRP1 causes 100-fold lower production of infectious virus than controls. Our results indicate that LRP1 is another DENV receptor thus, becoming an attractive target to evaluate for the development of effective antiviral drugs against DENV.

**Author summary:** Dengue virus (DENV) is a complex of four related viruses, recognized as serotypes, designated as DENV1-4. Any of the four DENV serotypes can cause a self-limited disease of mild flu-like symptoms known as dengue or its life threatening form, severe dengue, with hemorrhagic manifestations, organ impairment and shock. This disease is widely spread in tropical and sub-tropical areas worldwide, where the incidence of severe dengue has been increasing steadily. So far, efforts that target components of the viral replication machinery in order to develop a specific antiviral drug for dengue disease patients have failed. Thus, identifying the cell surface receptors used by DENV to enter host cells would provide a new molecular target to develop inhibitory drugs. Here, we evaluate the Low density lipoprotein receptor-related protein-1 (LRP1) as a putative DENV receptor. We present evidence demonstrating that LRP1 binds DENV through the viral envelope protein. We show that the production of infective virus is impaired on cells lacking LRP1, and that purified LRP1 is a potent blocker of DENV infection. These results are consistent with LRP1 playing an important role on DENV entry, making this receptor a molecule of interest on the investigation for medical treatments of dengue/severe dengue disease.

## Introduction

The Dengue complex consists of four viral serotypes (DENV1-4) of a mosquito-borne virus of the family *Flaviviridae*; genus Flavivirus. DENV-caused disease encompasses a wide spectrum of clinical manifestations ranging from a mild self-limited disease known as Dengue Fever to the life-threatening manifestations of Dengue Hemorrhagic Fever [1]. The global incidence of DENV infections has grown dramatically in recent decades, making this an arbovirus of worldwide public health concern [2].

DENV particles consist of a nucleoprotein core formed by RNA and the capsid protein. This core is surrounded by a host derived lipid bilayer with two glycoproteins inserted: the membrane (M) protein or its precursor (prM), found in immature virions and the envelope (E) protein [3]. This latter is organized in three structural domains DI, DII and DIII [4]. Of these, the DIII has been identified as directly involved on virus:receptor interactions [5–8]. However, no specific receptor has been identified that binds the virus through this region of the E protein.

DENV enters cell by receptor-mediated endocytosis [9]. Early steps of DENV infection comprise interactions with soluble and membrane-bound attachment factors, signaling receptors and internalization receptors [10].There are two main scenarios for these events, the primary infection with any of the four virus serotypes and a secondary heterologous infection [10]. In the latter, antibodies generated during the first infection cross-reacting but not cross-neutralizing the infecting serotype act as soluble attachment factors that bridge the virus to Fc□ receptors in the cell surface. Interaction with Fc□ receptors also appears to facilitate virus internalization [11, 12] and to generate intracellular signaling that suppress intracellular antiviral responses [13, 14]. All these factors together lead to a significant increase in the number of infected cells and virus production. This phenomenon, known as antibody dependent enhancement, results in an increased susceptibility of infected individuals to the development of severe forms of DENV-caused disease.

DENV also bind proteins such as Apolipoprotein A1 (ApoA1) [15] and Growth arrest-specific protein 6 [16] that bridge the virus with cell surface receptors, the scavenger receptor class B type I (SR-BI) and Tyrosine-protein kinase receptors Tyro 3, AXL and MER (TAM), respectively. SR-BI and TAM receptors have been shown to increase virus production but their role on early steps of infection has not been completely elucidated. Other well characterized attachment factors that DENV binds on the cell surface are heparan sulfate glycosaminoglycans [17], the C-type lectins DC-SIGN [18, 19] and L-SIGN [20] and the mannose receptor [21]. DC-SIGN and the mannose receptor have also been shown to act cooperatively in DENV binding to the signaling receptor C-type lectin domain family 5 member A (CLEC5A) [22]. This interaction enhances human macrophage CLEC5A/DAP12 signaling as part of the innate immune response. Thus, complexes formed by DENV attachment factors with signaling receptors may function in favor or against the infection.

Internalization receptors mediate virus entry via receptor-mediated endocytosis which ultimately leads to the delivery of the viral genome into the cytoplasm [23]. So far, only the T-cell immunoglobulin and mucin domain 1 (TIM-1) receptor, which binds phosphatidylserine moieties on the surface of the viral membrane, has been convincingly shown to mediate DENV endocytosis [24]. However, some cellular models that do not express TIM-1 are still sensitive, at low levels, to DENV infection. Therefore, we have not reached yet a complete understanding of the process whereby DENV gains entry into mammalian and mosquito cells, and additional molecules, binding either the phospholipids of the viral surface or the E protein, may participate in receptor-mediated endocytosis of this pathogen.

Regulation of lipid metabolism influences significantly the efficiency of DENV infection and replication *in vitro* e *in vivo* [25–28]. Sequence stretches on DENV E and capsid proteins resemble receptor binding motifs of ApoE [29]. ApoE is present in many lipoproteins and interacts with several receptors of the low-density lipoprotein receptor (LDLR) family [30]. Bovine lactoferrin, partially inhibits DENV and Japanese encephalitis infection presumably competing for binding to the LDLR [31,32]. Collectively, these results have suggested that LDLR may be involved on DENV entry to target cells. Yet, conclusive evidences of the participation of LDLR on DENV attachment and/or entry have not been obtained.

The LDLR family comprises at least 13 separate cell receptors that mediate lipoprotein internalization, although some also play a role in the regulation of cellular physiology and intracellular signaling [33]. In the context of the early events of the DENV infectious cycle we decided to focus our interest on a different member of the family, the LRP1 receptor. This protein plays a role not only in lipoprotein endocytosis but also in the internalization of protease:serpin complexes and, particularly of activated alpha-2-macroglobulin (α2M*):protease complexes [34]. We have found that proteins participating in these processes are significantly enriched within the plasma DENV interactome [35,36]. Also we have shown that α2M*, a ligand of LRP1, directly binds DENV virions and favors DENV infection [37].

## Materials and Methods

### Cells and viruses

Human hepatoma derived cells (Huh7) and human embryonic kidney cells (HEK-293T) cells were grown under standard conditions at 37°C, 5% CO_2_ in Dulbecco’s modified Eagle’s medium (DMEM) + GlutaMAX supplemented with 10% of fetal bovine serum (FBS), adding heat-inactivated FBS to a final concentration of 10% (v/v).

### Proteins

Recombinant proteins DVE1 (code MBS144575), DVE3 (code MBS 144886) and DVE4 (code MBS 145224) were purchased from MyBioSource, Inc, USA. The procedures for the expression, purification and general characterization of the proteins corresponding to the domain III of the E protein of the four DENV serotypes namely DIIIE1-4, were conducted as previously described. [37]. Receptor-activated α2M was purified from human plasma also following the procedure described elsewhere [37].

The human protein associated to the LRP1 receptor, known as RAP was obtained by expressing in *E. coli* a gene fragment coding for this molecule. The gene for RAP protein was amplified from the cDNA by PCR using the oligonucleotides **CATATG**TACTCGCGGGAGAAGAACCAG and **CTCGAG**TCAGAGTTCGTTGTGC, bearing on their 5’ end the recognition sequence for the restriction enzymes Nde I and Xho I (in boldface in the sequence). The amplified fragment was used to construct pET-RAP, a plasmid that codes for the intracellular synthesis of RAP under control of the T7 promoter. For protein expression, pET-RAP plasmid was transformed into the *E. coli* strain BL21. Recombinant RAP protein was obtained with a tag of 6 consecutive histidines at the N-terminus. RAP protein was purified from the supernatant by immobilized metal affinity chromatography using Ni-NTA agarose and eluted with a linear gradient of 10 to 300 mM imidazole in phosphate buffered saline (PBS, 10 mM Na_2_HPO_4_, 1.8 mM KH_2_PO_4_, 2.7 mM KCl, 137 mM NaCL, pH 7.4).

Recombinant proteins sLRP1_CII, sLRP1_CIII and sLRP_CIV were purchased from R&D Systems, USA. The Flavivirus-reactive mAb 4G2 was provided by the Unit for Production of Monoclonal Antibodies at the Center for Genetic Engineering and Biotechnology in Sancti Spiritus, Cuba. Mab MCA1965, was purchased from AbD serotec, USA and Mab 5A6 was donated by Dr. D. K. Strickland.

### Affinity purification of LRP1

Frozen human plasma was obtained from local blood banks. Pooled human plasma was dialyzed against 50 mM Hepes pH 6, 60 mM NaCl, 1 mM EDTA, centrifuged at 10,000 x g for 30 min at 4°C and filtered through 0.45 μm membrane. The resulting sample was fractionated using anion exchange chromatography as previously described [38]. Briefly, conditioned plasma was loaded onto a chromatography column packed with DE-52 gel previously equilibrated with the same buffer used for the dialysis. The sample eluted by step-washing of the column with 50 mM Hepes pH 6, 1 mM EDTA and increasing concentration of NaCL (0.3M, 6M and 1M) was then conditioned for loading in the affinity chromatography using receptor-activated α2M as immobilized ligand. Pooled elution fractions were dialyzed against 50 mM HEPES pH 7.0, 120 mM NaCl, 5 mM CaCl_2_, 1 mM MgCl_2_ and supplemented with a protease inhibitor cocktail (1 μg/ml leupeptin, 1 μg/ml pepstatin A, 1 μg/ml soybean trypsin inhibitor and 1 mM PMSF). Then, sample was loaded into a column packed with the affinity matrix at a flow of 10 cm/h. The column was washed with 100 column volumes of the loading buffer. Bound protein was eluted with 50 mM HEPES pH 6.0, 1.2 M NaCl, 25 mM EDTA. Eluted protein sample was dialyzed against the loading buffer, filtered through 0.2 μm and stored at −20°C until used.

### Surface plasmon resonance analysis (SPR)

SPR analyses were performed on a Biacore X unit (General Electric, USA). Proteins were covalently immobilized on CM5 chips using amine coupling chemistry as previously described [37]. DIIIE1 (30 μg/ml) and RAP (20 μg/ml) proteins were dissolved in 10 mM sodium acetate buffer pH 5 and loaded into the system. Immobilization of sLRP1_CIV was carried out at 10 μg/ml in 10 mM sodium acetate buffer pH 4.0. All SPR experiments were performed at a flow rate of 25 µl/min using Hepes 10 mM, pH 7.4, 0.15 M NaCl, 3 mM CaCl_2_, 0.005% surfactant P20 (HBS-Ca) as running buffer. The interacting surface was regenerated with a pulse of 5 µl of 10 mM NaOH. Results were analyzed using the BIAevaluation ver. 4.1 software application package (General Electric, USA).

### Pull down assays

Purified DIIIE1-4 and RAP were immobilized at a ligand density of 4 mg/mL in CNBr-activated Sepharose (Amersham, UK), as recommended by the provider. A background control matrix was prepared following the identical procedure but without any immobilized protein.

To obtain the microsomal fraction from Huh7 cells, subconfluent monolayers were scraped from the flasks, washed three times with PBS, resupended in 1 mL of hypotonic buffer (20 mM Tris pH 7.4, 10 mM KCl, 5 mM EDTA, 1 mM PMSF, 1 μg/ml peptstatin, 1 mM leupeptin) and incubated 20 min on ice. All subsequent steps were performed at 4°C. Sucrose was added to a final concentration of 250 mM, the cell suspension was transferred to a Dounce homogenizer and disrupted by 3 strokes followed by a brief sonication. Cell homogenates were first centrifuged at 1 000 x g to remove nuclei and supernatants were then centrifuged at 10 000 x g for 20 min to pellet mitochondrial fraction. The microsomal fraction was obtained by centrifuging post-mitochondrial supernatant at 100 000 x g for 1 h. The resulting 100 000 g pellet was resuspended at 10 mg/ml in HBS-Ca and stored at −80 °C until use.

The microsomal fraction was split in 6 identical aliquots, each of which was incubated with 1 mL of the affinity matrices and background control gel for 18 h at 4°C in a rotary mixer. The affinity gels were washed with 100 matrix volumes of HBS-Ca, 0.005% n-Octyl-β-D-Glucopyranoside. Bound proteins were eluted by incubating the gels twice for 15 min at 25°C with 2 mL of 0.1 M acetic acid followed by 1 mL of 3 M urea, 1 mM EDTA. The three eluates from each gel were then pooled, concentrated and adjusted to pH 8.0 by adding 0.2 mL of Tris-HCl 2 M, pH 8.0. Protein digestion, Selected Reaction Monitoring analysis and data processing were performed as described [35].

### Virus assays

Both viruses DENV2 (strain S16803, NIBSC code S16803) and Vesicular Stomatitis Virus (VSV strain Indiana) were propagated and titrated by plaque formation assays performed in Vero cells. Plaque formation assays were performed in 24 well plates with cell monolayers at 90% of confluence. Serial dilutions of virus inoculum were diluted in DMEM 2 % FBS, added to cells and incubated for 2 h in the CO_2_ incubator. Next, viral dilutions were removed, cell monolayers were washed twice with DMEM 2% FBS and incubated for 5 days at 37°C in high density medium (MEM supplemented with non-essential amino acids, 1% FBS, 1% carboxymethylcellulose) in order to propitiate the appearance of lytic plaques. The plaques were visualized by staining with 0.1% Naphtol Blue Black in 0.15 M sodium acetate. Two replicates were used per experimental point in each assay.

For the purification of DENV2, monolayers of Vero cells were infected at a moi 0.01, wash with fresh DMEM without FBS and maintained in serum free medium for 96 hrs. Cell supernatants were clarified by centrifugation at 5000 rpm for 30 min. Virus were sedimented by ultracentrifugation at sucrose 30% in 5 mM HEPES, 150 mM NaCl, 0.1 mM EDTA at 32 000 rpm for 9 hours. Virus pellets were resuspended in the same buffer without sucrose. The purified virus preparation was analyzed by SDS-PAGE followed by Coomasie blue staining. Visible protein bands corresponded to E and pre-M proteins as confirmed by western blotting using specific Mabs for each protein.

Inhibition of infection with chromatography fractions were performed using a plaque formation assay in Huh7 cells. Cells were seeded at 10^4^ per well in 24-well plates and kept for 18 h at 37°C in a CO_2_ incubator. DENV2 inoculum was diluted in DMEM without FBS and adjusted to obtain 50 PFU/well. Chromatography fractions were diluted in DMEM without FBS, added to the virus inoculum to obtain a final protein concentration of 10 μg/ml and incubated for 30 min at 37°C. For virus control, only PBS was added to the virus inoculum previous to the incubation. Next, virus-sample mixtures were added to cell monolayers and infection proceeded for 2 h at 37°C. Afterwards, cell monolayers were washed and high density medium was added. Viral plaques were visualized 5 days post-infection.

For the evaluation of virus yield after transfection with shRNAs, Vero cells were seeded in 96 well plates and cultured until monolayers reached 90% of confluence. Viruses were diluted in DMEM, 2% FBS for an m.o.i of 0.1 in case of DENV2 and 0.35 for VSV. Monolayers were washed once with DMEM 2% FBS and subsequently 100 µl of virus inoculum were added. After 1 h incubation viral inoculum was removed and replaced by 100 µl of fresh DMEM 2% FBS. To evaluate viral production, 24 h supernatant was collected and assayed for infectious virus by plaque assay on Vero cells.

### Gene silencing

Target sequence within the hLRP1 mRNA (5’-GATCCGTGTGAACCGCTTTAA-3’) (NM_002332.2, RNAi Consortium at http://www.broadinstitute.org/rnai/public) was confirmed to match within a CDS on the LRP1 gene sequence and to be unique for this gene. Template sequences encoding a short hairpin RNA were generated and cloned into de pKLO.1-TRC shRNA lentivector:

Forward oligo: 5’-CCGGGATCCGTGTGAACCGCTTTAACTGCAGTTAAAGCGGTTCACACGGATCTTTTTG-3’
Reverse oligo 5’-AATTCAAAAAGATCCGTGTGAACCGCTTTAACTGCAGTTAAAGCGGTTCACACGGATC-3’

Oligonucleotides (Integrated DNA Technologies, USA) were annealed, phosphorylated and ligated into AgeI and EcoRI sites of the pKLO.1 vector. Plasmid containing clones were selected by resistance to ampicillin and checked by restriction analysis and sequencing.

Monolayers of 293T cells were co-transfected with packaging construct (psPAX2) coding for gag, pol, tat and rev of HIV; envelope plasmid (VSV-G) and pLKO.1 based shRNA-LRP1. As negative controls were used two pLKO.1 constructs designed to activate the silencing machinery but without targeting any gene in the cells: shRNA-scramble. Transfection was performed in 10 cm plates by the Ca-phosphate method using 10 µg of DNA mixed in a ratio pLKO.1_shRNA:psPAX:VSV-G 4:3:1. After 24 hours, media containing transfection mixtures was replace by fresh DMEM, 10% FBS. Lentivector containing supernatants were harvested at 48 and 72 hours post-transfection. Collected supernatants were clarified by centrifugation followed by filtration through 0.45 μm, aliquoted and store at −80°C until used.

Lentivirus containing supernatants were used at indicated dilutions to infect Huh7 cells. Infection was performed in DMEM, 10% FBS, 5 μg/ml polybrene. After 18 h lentivirus inoculum was removed and replaced by selection medium: DMEM, 10% FBS, 1 μg/ml Puromycin (InvivoGen, USA). Efficiency of transduction was estimated by detection of GFP expression using FACS.

Huh7 cells transduced with either the LRP1-specific shRNA or a scrambled shRNA control were infected at day 7 post-transduction with DENV2 and infectious particles release was quantified by titration 24 h post-infection.

### Denaturing polyacrilamide gel electrophoresis (SDS-PAGE)

One microgram of protein of chromatography fractions was diluted in non-reducing sample buffer, loaded in a 6−15% gradient polyacrylamide denaturing gel and separated under standard conditions [39]. Proteins were detected using silver staining [40]. For the evaluation of LRP1 expression after gene silencing 10^5^ cells were lysed in non-reducing sample buffer, separated in an 8% polyacrylamide gel and subjected to western blotting analysis.

### Western blotting

Separated proteins were transferred to nitrocellulose membranes under standard conditions [41]. After transfer membranes were incubated for 1 hr at 25°C with Hepes 20 mM pH 7.2, 0.05% tween 20, 1 mM CaCl_2_. Membranes were then blocked for 1h using 1% Bovine Serum Albumin (BSA) and incubated with a 10 μg/ml dilution of primary antibody for 16 hrs at 4°C. Next, blots were washed three times with PBS-T and antibody binding was detected using an anti-mouse antibody conjugated to HRP. For visualization, ECL development solution was used in the evaluation of LRP1 expression after gene silencing and 3,3’-diaminobenzidine (Sigma, USA) in the analysis of affinity chromatography fractions.

### ELISA

Ninety-six well microtiter plates were coated for 1 h at 37°C with 50 μL per well of the corresponding protein at 10 μg/ml or 2×10^6^ pfu of purified DENV2 in carbonate buffer pH 9.6. The plates were blocked with 2% (w/v) BSA in PBS, 0.05% Tween 20 (PBS-T) for 1 h at 37°C. Recombinant sLRP-CI-IV were diluted in HBS-Ca, added to coated wells and incubated 1 h at 37°C. After three washes with binding buffer, bound proteins were detected using an anti-human Ig, Fc specific-HRP conjugate. The development was carried out with 1 mg/ml o-phenylenediamine dihydrochloride, 0.1% hydrogen peroxide in 0.05 M phosphate-citrate buffer, pH 5.0. Developing reaction was stopped with 3 M H_2_SO_4_ and results were read at 492 nm in a microplate reader.

### Statistical Analysis

GraphPad Prism V5.3 was used to perform unpaired Student’s t-test to calculate p values [42]. Differences with P-values of <0.05 were considered significant (*p<0.05, **p<0.01, ***p <0.001).

## Results

### LRP1 depletion inhibits DENV2 infection

To investigate whether the presence of LRP1 on the cell surface could impact the efficiency of infection, we examined the effect of sh-RNA mediated LRP1 knock-down on viral yield. The assay was performed in Huh7 hepatoma cells, which are extensively used in *Flaviviridae* research, physiologically relevant for DENV infection [43] and express LRP1 [44]. Huh7 cells can be efficiently transduced by lentiviral vectors without alterations to key hallmarks of the hepatic phenotype [45]. Western blotting analysis revealed that LRP1 expression was efficiently inhibited by day 6 post-transduction (Fig 1A). Viral yield in the cells transduced with the LRP1-specific shRNA was 100-fold lower than in the cells transduced with the scrambled control (Fig 1B). These results show that LRP1 plays a central role on DENV replication cycle. Meanwhile, VSV which presumably can use LRP1 as an alternative to LDLR [46], significantly less efficient than this latter, exhibits only a slight decrease in virus titer.

**Fig 1.**
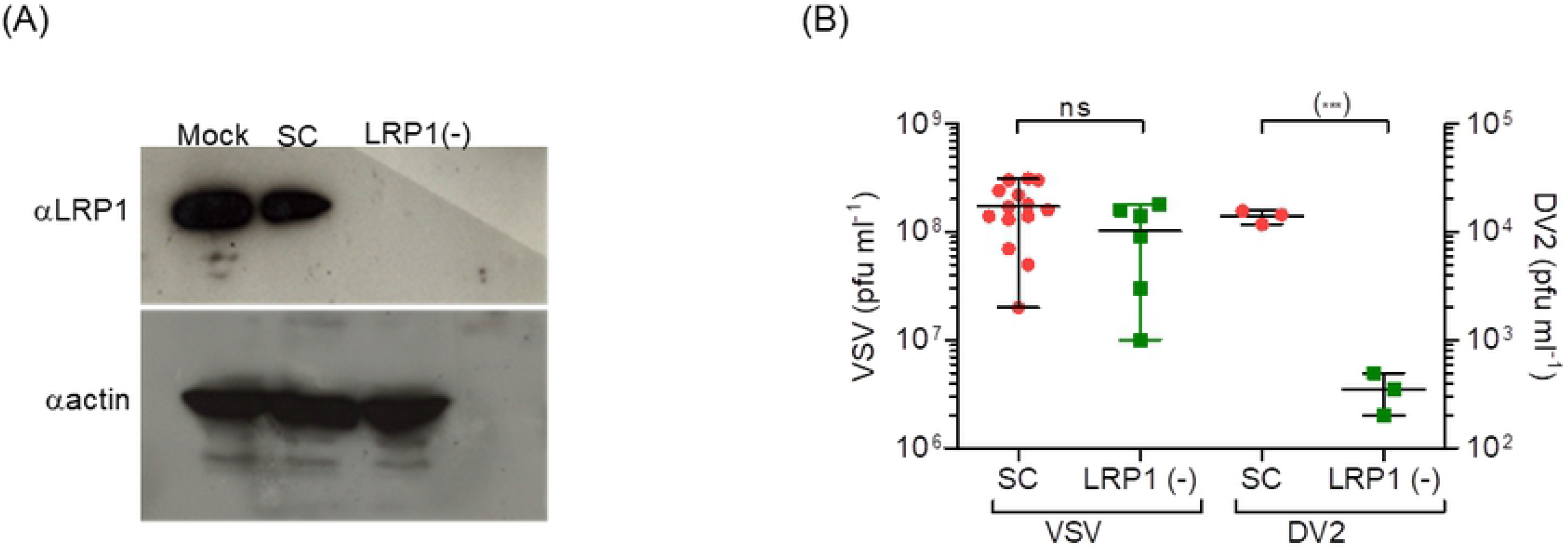
LRP1 is essential for DENV2 infection. (A) Western blotting analysis of the silencing of LRP1 expression on cells transfected with scramble shRNA (CS) or with LRP1-shRNA. LRP1 was detected using the Mab 5A6 that recognizes the 85-kDa β chain of the receptor [47]. Detection of βactin was performed on the same membrane. (B) Effect of the transduction of LRP1-shRNA on DENV2 infection. Transduced cells were infected at day 7 with DENV2 (moi of 0.1) or Vesicular Stomatitis Virus (VSV, moi 0.35). Virus yield was measured on 24 hr post-infection supernatants.

### Ligand-purified LRP1 inhibits DENV2 infection in hepatocytes

One testable prediction of the hypothesis that LRP1 participates on early steps of virus:cell interaction would be to inhibit DENV infection by competition with a soluble form of LRP1. Therefore, we purified the LRP1 ectodomain (sLRP1) from human plasma by using receptor-activated α2M as an affinity ligand (Figs 2A-C), taking advantage of the fact that under physiological conditions sLRP1 is shed from cell surfaces into body fluids in a soluble, active ligand-binding form [48]. SPR experiments with the purified sLRP1 on chips containing immobilized RAP, which is the intracellular receptor chaperone of LRP1 [49], confirmed that it retained its normal ligand-binding activity (Fig 2D).

**Fig 2.**
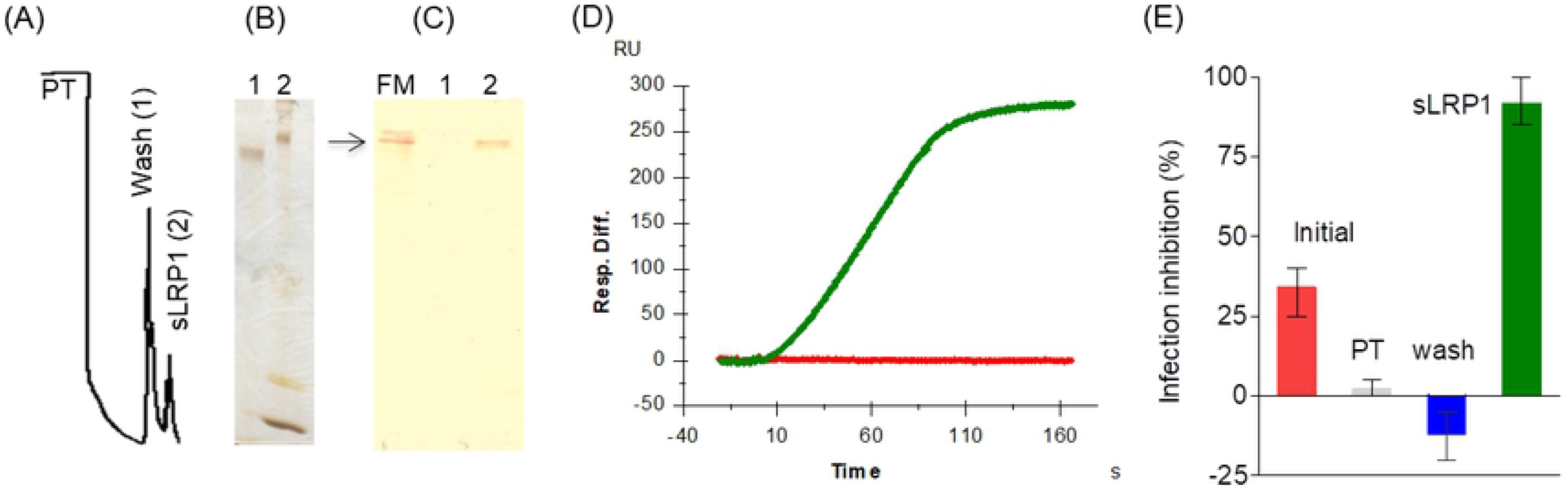
Purified sLRP1 inhibits DENV2 infection. (A) Chromatographic profile of the affinity chromatography using receptor-activated α2M as immobilized ligand. PT: pass-through. Numbers in parenthesis identify the lane in SDS-PAGE (B) and western blotting analysis. In (C) membrane was probed with an anti-LRP1 Mab (MCA1965) that recognizes the alpha chain of the receptor. (C). Microsomal fraction (FM) of Huh7 cells was used as control of migration of α chain of LRP1. (D) Binding analysis of purified sLRP1 to recombinant RAP protein using SPR. RAP protein was immobilized on the surface of the chip and purified sLRP1 was injected diluted in HBS-Ca at 1 μg/ml. (E) Inhibition of DENV2 infection in Huh7 cells after incubation with chromatography fractions. Results from one representative experiment out of three independent experiments. Error bars represent the range of three sample replicates.

An inhibition-of-infection assay was then set up in Huh7 cells where a DENV2 preparation was pre-incubated either with purified sLRP1 or with different fractions collected during the purification process (the initial sample, the pass-through and the washing eluate). As previously reported, on these experimental conditions the human plasma fraction used as starting sample shows partial inhibitory activity against DENV2 infection of Huh7 hepatocytes [50]. Neither the pass-through nor the washing eluate exhibited inhibitory activity, while the eluate containing sLRP1 at 5×10^−8^ mol/L, exhibited an inhibitory activity close to 90% indicating a significant enrichment of inhibitory activity (Fig 2E). These results indicate a direct interaction of LRP1 with DENV2 and are consistent with the involvement of LRP1 in the initial stages of the DENV life cycle. The high potency of the observed inhibitory effect suggests that it involves the blockage of critical functional sites on the viral surface (*i.e.* direct competition with cell-bound LRP1) rather than the indirect occlusion of binding sites for other cell receptors.

### LRP1 interacts with DENV particles

LRP1 is a heterodimeric type I integral membrane protein comprised of an extracellular 515-kDa α chain and a non-covalently linked cytoplasmic and transmembrane 85-kDa β chain (Fig 3A). The interaction of LRP1 with extracellular ligands typically takes place through the complement-type repeats that are individually known as ligand binding domains (LBD) and are presented in four clusters of 2, 8, 10 and 11 LBD on the α chain denominated CI, CII, CIII and CIV, respectively. Of these, clusters CII-CIV are those that bind most of LRP1 ligands [49].

**Fig 3.**
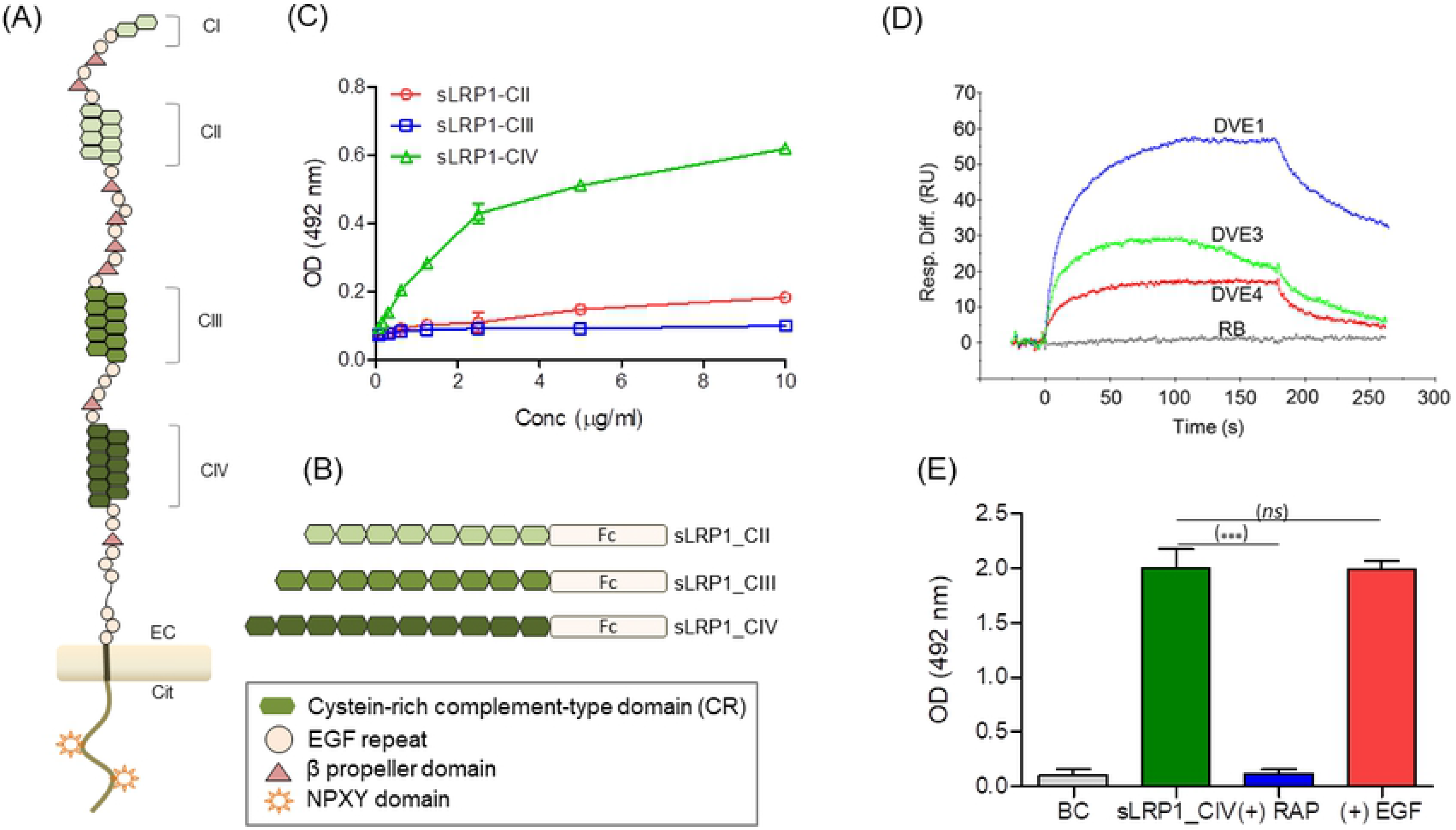
Interaction of DENV2 with ligand binding clusters of LRP1. Schematic representation of (A) LRP1 and (B) the recombinant proteins sLRP1-CII, sLRP1-CIII and sLRP1-CIV. (C) Evaluation of the interaction of recombinant proteins sLRP1-CII, sLRP-CIII and sLRP1-CIV with purified DENV2. (D) Interaction of sLRP1-CIV with recombinant preparations of the ectodomain of the E protein of DENV serotypes 1, 3 and 4 (DENVE1, DENVE3 and DENVE4) by SPR. sLRP1-CIV was immobilized in the surface of the chip and recombinant DENV E proteins were injected diluted in HBS-Ca at 10 μg/ml. Resp Diff. Signal obtained by online subtraction of background control channel. (E) Binding competition by ELISA. Plates were coated with DENVE1 protein and the binding of sLRP1-CIV at 5 μg/ml was evaluated after the incubation for 30 min at 37°C with EGF or the LRP1 ligand protein RAP, both proteins used at 5 μg/ml. BC: background control. Results are representative of two independent analyses. Error bars represent the range of three replicates.

In order to investigate whether DENV particles can bind LRP1, we analyzed the interaction of a purified preparation of DENV2 adsorbed to ELISA plates with three chimeric proteins denoted sLRP1_CII, sLRP1_CIII and sLRP1_CIV, containing clusters II, III or IV from LRP1 fused to a human IgG1 Fc domain (Fig 3B). As shown in Fig 3C, DENV2 bound efficiently the fusion protein containing cluster IV, but not those containing clusters II or III. Considering that the construct of sLRP1-CIV protein comprises only the 11 LBD of CIV (Fig 3B), these results directly imply that, similar to the interaction with other extracellular ligands of LRP1, the complement-type repeats (Fig 3A) are involved in the interaction with DENV2.

### The E protein of DENV binds LRP1

The E protein covers most of the solvent-accessible surface of DENV virions [51]. Therefore, we next investigated if protein E mediates the interaction between DENV and LRP1 and whether this interaction is conserved for the other viral serotypes. To this end, we examined by SPR the binding of recombinant proteins encompassing the E ectodomain from DENV serotypes 1, 3 and 4 (DVE1, DVE 3 and DVE 4 in Fig 3D) to sLRP1_CIV immobilized on an SPR chip. All three recombinant E proteins bound the sLRP1_CIV-derivatized surface obtaining almost three times higher signal for DVE1 followed by DVE3 and DVE4 (Fig 3D). This result demonstrate that the DENV-LRP1 cluster IV interaction is mediated by protein E and that it is preserved in serotypes other than DENV2, although with probable differences in reactivity among the four viral serotypes. DVE1 was then selected to test whether RAP protein was able to block the interaction using an ELISA format. Results showed that similar to many other LRP1 ligands, pre-incubation of sLRP1-CIV with RAP abolished the interaction with DVE1 while no effect was observed when another protein non-relevant for this system *i.e*. the human Epidermal Growth Factor (EGF), was used as competitor (Fig 3E).

### The interaction of E with LRP1 is mediated by DIII of protein E

A number of results indicate that DIII of E protein of Flaviviruses participate on the interaction with cell surface molecules that play key roles in virus binding [5,7,8,52,53]. We thus decided to investigate whether the protein E-LRP1 interaction is mediated by this region of E protein. For this purpose, we used recombinant DIII proteins of the four viral serotypes, denoted DIIIE1, DIIIE2, DIIIE3 and DIIIE4 (residues 289-400 of DENV1, 2, 4 and 287-388 of DENV3) (Fig 4A), to coat ELISA plates, which were probed with the sLRP1-derived chimeras (Fig 3B). DIIIE from all four serotypes bound sLRP1-CIV (Fig 4B). DIIIE2 also bound sLRP1_CII weakly, although this reactivity was undetectable for the other three serotypes, and no binding to sLRP1_CIII was detected for any of the recombinant DIII preparations. In this assay format, the recombinant protein corresponding to DIII of DENV1 (DIIIE1) was the one showing the lowest reactivity. This apparent contradiction with the results obtained by SPR with DENVE1, 3 and 4 (Fig 3D) may reflect a higher susceptibility of the recombinant DIIIE1 protein to loose reactivity when adsorbed to the surface of the plate. In fact, DIIIE1 showed similar reactivity with purified sLRP1 as DIIIE2 when compared in SPR (Fig4C). Specificity of the binding to DIIIE1 surface was evidenced using competition assay with two LRP1 ligands, RAP and receptor activated α2M. As shown in Fig 4C binding to DIIIE1 surface decreases significantly when RAP and receptor activated α2M were incubated with sLRP1 before the analysis.

**Fig 4.**
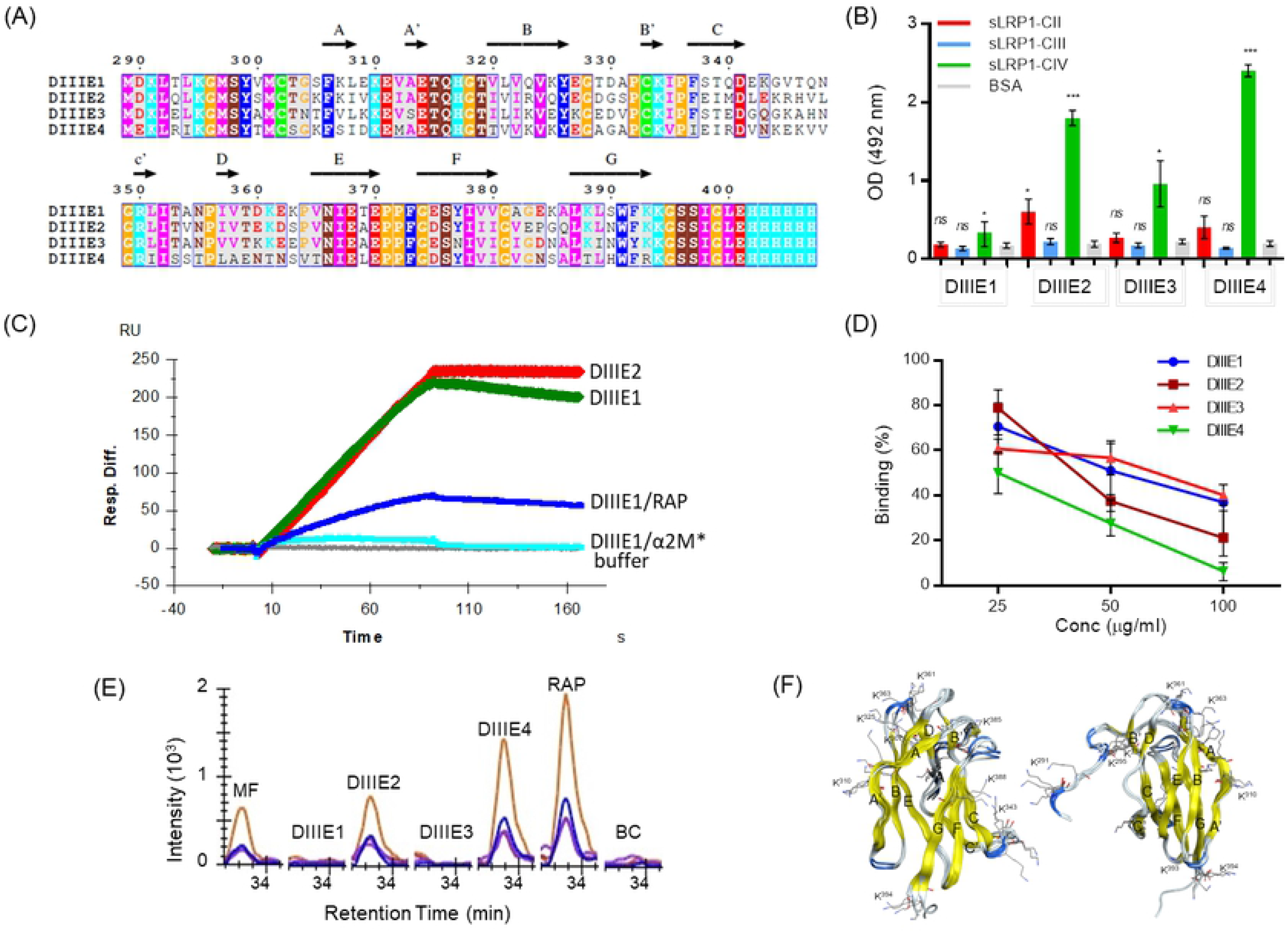
Interaction of recombinant DENV E DIII proteins with LRP1. (A) Multiple sequence alignment of DIIIE1-4 proteins (residue numbers according to DENV1, 2 and 4). The alignment was performed using the ClustalW application [54]. The arrows denote β-strands. Residues are colored according to the physico-chemical properties and conservation according to ESPript program [55]. (B) Interaction of DIIIE1-4 proteins with sLRP1-CII, sLRP1-CIII or sLRP1-CIV by ELISA. Results are mean and SEM of three independent experiments. (C) Binding analysis of sLRP1 to immobilized DIIIE1 by SPR. sLRP1 at 1 μg/ml was loaded over the DIIIE2 and DIIIE1 surfaces. For this later, sLRP1 injection was performed without or after pre-incubation with RAP at (DIIIE1/RAP), or receptor-activated α2M (DIIIE1/α2M*). Resp. Diff.: non-specific signal was online subtracted using a reference channel without any immobilized protein. (D) Blocking of sLRP1-DENV2 binding. sLRP1-CIV at 5 μg/ml was incubated with 10 μg/ml of DIIIE1-4 for 1 h before addition to DENV2 coated plates. Bound sLRP1-CIV was detected using an anti-human Fc Ig-POD conjugate. (E) Binding to LRP1 present in the microsomal fraction of Huh7 cells. Pull down assay using recombinant DIIIE1-4 proteins as baits and detection by Mass spectrometry using Selected Reaction Monitoring. MF: microsomal fraction. BC: background control. Data shows transition signals of the surrogate peptide _365_IVFPHGITLDLVSR_378_ (Swissprot Accession number Q07954). Samples were analyzed in duplicate. Similar results were obtained for other five proteotypic peptides (S1 Table). (F) Cartoon representation of the 3D structure of DIII. In yellow are represented the β-strands and lysine residues are shown in wire-frame representation. Residue numbers correspond to DENV1, 2 and 4.

The ability of recombinant DIII proteins to block the binding of sLRP1-CIV to DENV2 (Fig 4D) was also evaluated. In the assay, DIIIE1-4 proteins were incubated in solution with sLRP1-CIV before its addition to DENV2 particles adsorbed in ELISA plate wells. The results show that DIII recombinant proteins of the different viral serotypes are able to block sLRP1-CIV to DENV2 to different levels with more than 95% of inhibition exhibited by DIIIE4 at the maximal protein concentration evaluated in the assay (100 μg/ml) followed by DIIIE2 > DIIIE1 > DIIIE3. This result shows that DIII is a dominant site of interaction, although it does not rule out the existence of other regions within the E protein that may contribute to the binding to sLRP1-CIV. This result also highlights the fact that DIII surface patches involved on the interaction with LRP1 are sufficiently similar that a recombinant DIII from one serotype can effectively compete for LRP1 binding to the virus particle of a different serotype.

The binding of recombinant DIII to full-length LRP1 was evaluated in pull-down assays (Fig 4E). To this end, DIIIE1-4 were used as baits and incubated with a microsomal fraction obtained from Huh7 cells, detecting bound LRP1 by selected reaction monitoring in mass spectrometry. In this assay setup, both DIIIE2 and DIIIE4 were able to pull down the native LRP1 receptor as presented in the cell surface (Fig 4E and Table A in S1).

Altogether, these results demonstrate that the four serotypes of DENV bind the LRP1 receptor and that DIII mediated interaction is probably primary in binding to cluster IV of ligand binding domains of this receptor. Even considering a possible influence of different assay formats, DIII mediated interaction is stronger with serotype 4 followed by serotype 2 and weaker with serotypes 3 and 1.

Similar to other members of the family of the LDLR, LRP1 binding to its ligands has been proposed to occur through the acidic necklace model: amino groups of lysine residues of the ligands are encircled in a tripartite salt bridge via three remaining oxygen atoms from the acidic residues forming the octahedral calcium cage of LBDs [56]. It seems that there are modest requirements for the coordination of lysine residues [57], but specificity may be influenced by other residues located in its vicinity showing positive/negative charge [58,59] or hydrophobic character [60]. High affinity binding seems to involve the interaction with two or more LBDs requiring at least two lysine residues located at an appropriated distance - 18-20 Å according to structural data – which allows the simultaneous binding of two consecutive LBDs [56]. Likewise, the relative contribution of a single lysine residue of a ligand protein to the interaction may be difficult to rationalize in terms of stereo-chemical properties and the overall binding affinity may be the result of the additive effect of various weak binding sites [61].

The DIII contains 10-13 lysine residues depending on the serotype, but the number of basic residues is counted by the sum of aspartic and glutamic acids resulting in the fact that at neutral pH the overall charge is predicted to be minimal for DENV1-DENV3 and only approximately +2 for DENV4 (Table B in S1). This observation highlights the relevance of specific interactions involving lysine residues of DIII which interact with the receptor according to the acidic necklace model. The overall positive electrostatic potential of DIII from DENV4 may also contribute favorably to the interaction and explains in part why this serotype displays the higher binding affinity. Five lysine residues (K291, K295, K310, K334 and K394) are strictly conserved – in sequence – among the serotypes but others are conserved topographically *i.e.* the mutation of one lysine is accompanied by a compensating mutation to lysine in an adjacent position in the 3D structure, e.g. the mutation K388 (DENV1) to T (DENV4) is compensated by the mutation G344 (DENV1) to K (DENV4). Furthermore, there are several interatomic distances between the lysine residues of DIII which are consistent with a putative interaction with two consecutive LBDs. Otherwise, the sequence identity among the DIII from the four serotypes varies between 52 and 71%, indicating that mutations close in sequence and/or 3D space to lysine residues would also affect the interaction and explain observed differences in binding affinity (Fig 2A and Table C in S1).

## Discussion

Viral tropism is strongly determined, to a great extent, by the specific host cell receptor(s) leveraged by a virus to infect a specific cell or tissue type [62]. Thus, the fact that DENV infects a dissimilar array of cell types including endothelial cells, fibroblasts, myeloid cells and lymphocytes [63] might be explained by the use of a single, ubiquitously expressed receptor or of several, varying cell receptors differentially expressed in distinct cell lineages.

A substantial number of different cell surface molecules have been shown to be involved in early stages of the DENV:cell interaction; most of these cases concern the first step of the cell entry process, that is, adhesion [64]. The higher efficiency of infection afforded by the ectopic expression of DENV adhesion molecules, such as DC-SIGN, illustrates the importance of this initial interaction for the viral life cycle [18].

However, the following step, which in DENV case is cell entry by receptor-mediated endocytosis, is as important as the first. In the case of human cells, receptor-mediated endocytosis places the virion at the right cellular location and under the best conditions for membrane fusion [23]. Although TIM-1 is the only molecule that has convincingly been shown to internalize DENV by receptor-mediated endocytosis [24], both the experimental results of the characterization of TIM-1 activity during endocytosis [16,24] and the expression profile of this protein suggest that the virus must leverage other as yet unknown molecules capable of receptor-mediated endocytosis.

For a cell receptor to be used as a viral port of entry, it must be able to bind the virus directly or through a bridging ligand. Our results show that LRP1 binds DENV virions (Fig 3C) and that this interaction competes, with potency at the nanomolar range, with DENV virion interactions involving other cell surface molecules (Fig 2E). Previous research has already shown that DENV virions can also bind LRP1 ligands [36,65,66], and in the case of α2M, it has been proven that this interaction augments infection [65]. Whether DENV infection may proceed through the formation of a ternary DENV:α2M:LRP1 complex is the subject of ongoing research, but we would like to point out that the pathogenicity of WNV, a flavivirus closely related to DENV, is severely reduced in knockout mice defective for the expression of α2M homologs [67]. Lastly, it is worth noting that the assembly of ternary complexes is not exclusive to multivalent LRP1 ligands such as DENV and α2M, but has been observed for other LRP1 ligands as well [68].

LRP1 is a highly efficient constitutive endocytic receptor mediating the internalization of over 40 different ligands, including viruses [49]. A direct interaction with this receptor leads, with high probability, to the internalization of the ligand. Consequently, our results demonstrating that the expression of LRP1 in human hepatocytes is a requisite for efficient DENV infection in these cells (Fig 1) indicates that LRP1-mediated cell entry must result in a productive infection event.

Severe dengue patients have been shown to exhibit reduced LRP1 expression levels [69]. Based on this result, a proposal was put forth to include LRP1, together with another 17 genes, into a molecular signature for the prognosis of dengue disease [69].

In light of our findings regarding the direct interaction of DENV virions with LRP1, this proposal reinforces the notion that LRP1 is directly linked to the process of DENV infection *in vivo* and the pathogenesis of dengue disease, although whether its involvement is related to the virus entry stage or extends to the well-known participation of this molecule in diverse signaling and regulation processes [49] remains to be determined.

LRP1 is well conserved evolutionarily [49]. Its *A. aegypti* ortholog (NCBI Reference Sequence: XP_021706493.1) exhibits 35% sequence identity with the human protein, which is even higher (51%) across ligand-binding cluster IV, whose interaction with DENV is strongest (Fig 3C, 4B). Were LRP1 to play a role in the entry of DENV into mosquito cells similar to that described here in human cells, it would position this receptor as a prime target for the development of antiviral strategies. However, in contrast with our results, silencing of LRP1 orthologs in mosquito cells actually increases the copy number of viral RNA in infected cells [28]. This paradoxical result, according to the data published by Tree *et al*., stems from the involvement of LRP1 in the regulation of intracellular cholesterol, whereby lower LRP1 levels would increase the amount of intracellular cholesterol and, therefore, facilitate viral replication [28]. Undoubtedly, the precise role of LRP1 in DENV infection of human and mosquito cells deserves the focus of further research efforts.

Our data prove that LRP1 binds DENV virions through protein E (Fig 3D and E), or more specifically through DIII of protein E (Fig 4), although they do not rule out the possible involvement of additional regions. DIII is the target of many highly potent neutralizing antibodies [70–72] that have been shown to disrupt and block early interaction events with cell surface molecules–presumably, the viral cell receptors [73]. This, together with the immunoglobulin-like fold of this domain and the capacity of recombinant DIII to bind the cell surface [74] and compete with virus binding [6], has led many to consider it a receptor-interacting region, despite the fact that the only receptors DIII has been shown to bind are cell surface glycosaminoglycans [75]. The results presented here indicate that the binding of protein E to LRP1 is conserved across all four DENV serotypes, although there are considerable inter-serotype differences regarding its affinity (Fig 4D).

It is also worth noting that several of the molecules described as DENV adhesion receptors are members of well-known LRP1 functional complexes. For instance, many LRP1 ligands use glycosaminoglycans as adhesion molecules [76], and proteins hsp90 [77], grp78 [78] and AXL [79], which have been separately identified as DENV adhesion receptors [16,80,81], act cooperatively with LRP1 in binding, entry and signaling processes.

LRP1 is ubiquitously expressed, including several tissues and cells types identified as relevant for DENV replication such as monocytes/macrophages, keratinocytes, myeloid cells, phagocytes, hepatocytes and endothelial cells [49,82]. Thus, we hypothesize that LPR1 might constitute one of the hitherto unidentified endocytic receptors exploited by DENV to gain entry into mammalian cells.

## Acknowledgments

The authors would like to acknowledge Dr. Felix Rey and Dr. Nolwen Jouvenet, both from Pasteur Institute, Paris, France, for their valuable comments during the preparation of the manuscript.

## Supporting information S1

**Table A. Factor of enrichment of LRP1 proteotypic peptides respect to the background control in the Pull Down experiments.** ^1^P-value calculated for the T-test analysis of each experimental condition means (DIIIE1, DIIIE2, DIIIE3 or DIIIE4) versus the background means of normalized intensity values. (*), denotes P-value below 0.05, (ns) no significative. The Skyline software was used to carry out the statistical analysis.

**Table B. Isoelectric point and charge calculations.**

**Table C. Sequence identity* and similarity of DIII from DENV1-4**. *values located over (under) the diagonal correspond to sequence identity (similarity).

